# Comparison of 12 molecular detection assays for SARS-CoV-2

**DOI:** 10.1101/2020.06.24.170332

**Authors:** Yasufumi Matsumura, Tsunehiro Shimizu, Taro Noguchi, Satoshi Nakano, Masaki Yamamoto, Miki Nagao

**Affiliations:** Department of Clinical Laboratory Medicine, Kyoto University Graduate School of Medicine, Kyoto, Japan; Department of Infectious Diseases, Kyoto City Hospital, Kyoto, Japan

## Abstract

Molecular testing for SARS-CoV-2 is the mainstay for accurate diagnosis of the infection, but the diagnostic performances of available assays have not been defined. We compared 12 molecular diagnostic assays, including 8 commercial kits using 155 respiratory samples (65 nasopharyngeal swabs, 45 oropharyngeal swabs, and 45 sputum) collected at 2 Japanese hospitals. Sixty-eight samples were positive for more than one assay and one genetic locus and were defined as true positive samples. All the assays showed a specificity of 100% (95% confidence interval, 95.8 to 100). The N2 assay kit of the US Centers for Disease Control and Prevention (CDC) and the N2 assay of the Japanese National Institute of Infectious Disease (NIID) were the most sensitive assays with 100% sensitivity (95% confidence interval, 94.7 to 100), followed by the CDC N1 kit, E assay by Corman, and NIID N2 assay multiplex with internal control reactions. These assays are reliable as first-line molecular assays in laboratories when combined with appropriate internal control reactions.

## Introduction

Accurate detection tests for severe acute respiratory syndrome coronavirus 2 (SARS-CoV-2) are important to combat the coronavirus disease 2019 (COVID-19) pandemic (1). Various molecular diagnostic assays have been developed and used worldwide (1-4), but the differences in their diagnostic performances remain poorly understood. In this study, we aimed to compare the performance of 12 molecular assays.

## Materials and Methods

### Clinical specimens

A total of 923 upper or lower respiratory tract samples (nasopharyngeal swabs and oropharyngeal swabs in viral transport media or sputum) were collected from 446 patients who were suspected to have COVID-19 between January and May 2020 at Kyoto University Hospital and Kyoto City Hospital. In this study, we included all 68 SARS-CoV-2-positive samples and 87 randomly selected negative samples from 107 patients.

### RNA extraction

The respiratory samples were prospectively stored at −80°C after stabilization by mixing an equal volume of DNA/RNA Shield (2X concentrate; Zymo Research, Irvine, CA). The thawed samples were centrifuged at 20,000×*g* for 2 min. RNA was extracted from 140 μL of the supernatant using the QIAamp Viral RNA Mini Kit (Qiagen, Hilden, Germany) with RNA extraction controls—5 μL of LightMix® Modular EAV RNA Extraction Control (EAV; Roche, Basel, Switzerland) or 10 μL of MS2 phage (Thermo Fisher Scientific, Waltham, MA, USA)—and eluted in a final volume of 60 μL.

### Molecular assays

Table 1 shows the molecular assays evaluated in this study. Real-time RT-PCRs were performed using N1, N2, and RNaseP (RP) internal control assays developed by the Centers for Disease Control and Prevention, USA (2019-nCoV CDC EUA kit (5), obtained from Integrated DNA Technologies, Coralville, Iowa, USA), N2 assay developed and distributed by the National Institute of Infectious Disease (NIID) in Japan (4) (with/without EAV), N and E assays developed by Charité in Germany (1) (Corman) with TaqPath™ 1-Step RT-qPCR Master Mix, CG (Thermo Fisher Scientific). We also tested the LightMix® Modular assays (Roche) for E, RdRP, and N genes multiplexed with EAV, the Real-Time Fluorescent RT-PCR kit for detecting 2019-nCoV (BGI Biotechnology, Wuhan, China), and the TaqPath™ COVID-19 Combo Kit (Thermo Fisher Scientific) according to the manufacturers’ instructions. The above reactions were performed using a LightCycler® 480 System II (Roche), and cycle threshold (Ct) values were determined by the second derivative maximum method, except for the CDC N1/N2 and TaqPath™ COVID-19 Combo Kit assays, which were performed using Applied Biosystems® 7500 Fast or QuantStudio5 Real-Time PCR Systems (Thermo Fisher Scientific) using a fixed threshold of 0.1. A loop-mediated isothermal amplification (LAMP) assay was performed using a Loopamp® SARS-CoV-2 detection kit and LoopampEXIA® real-time turbidimeter (Eiken Chemical, Tokyo, Japan).

**Table 1.**
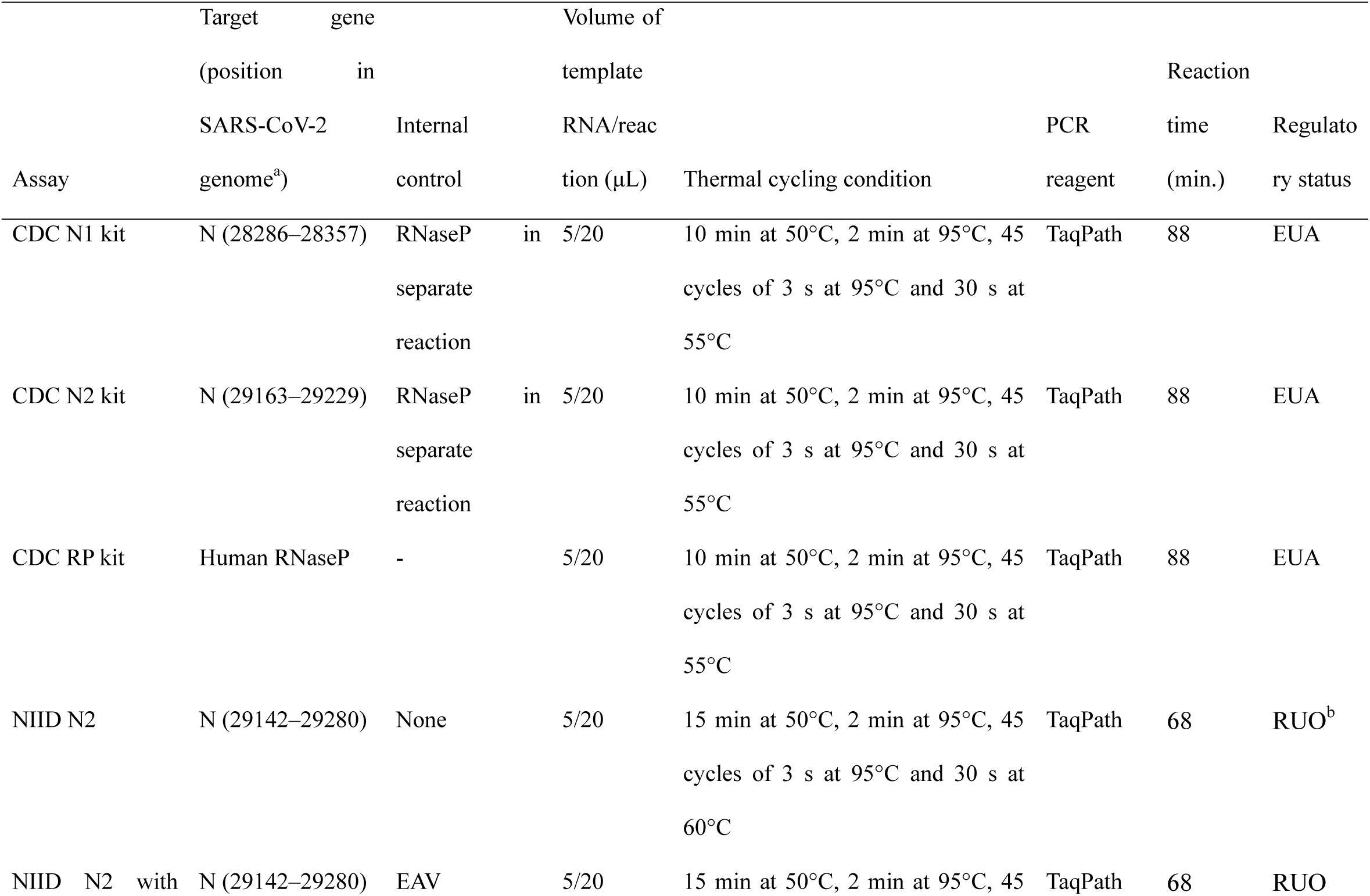
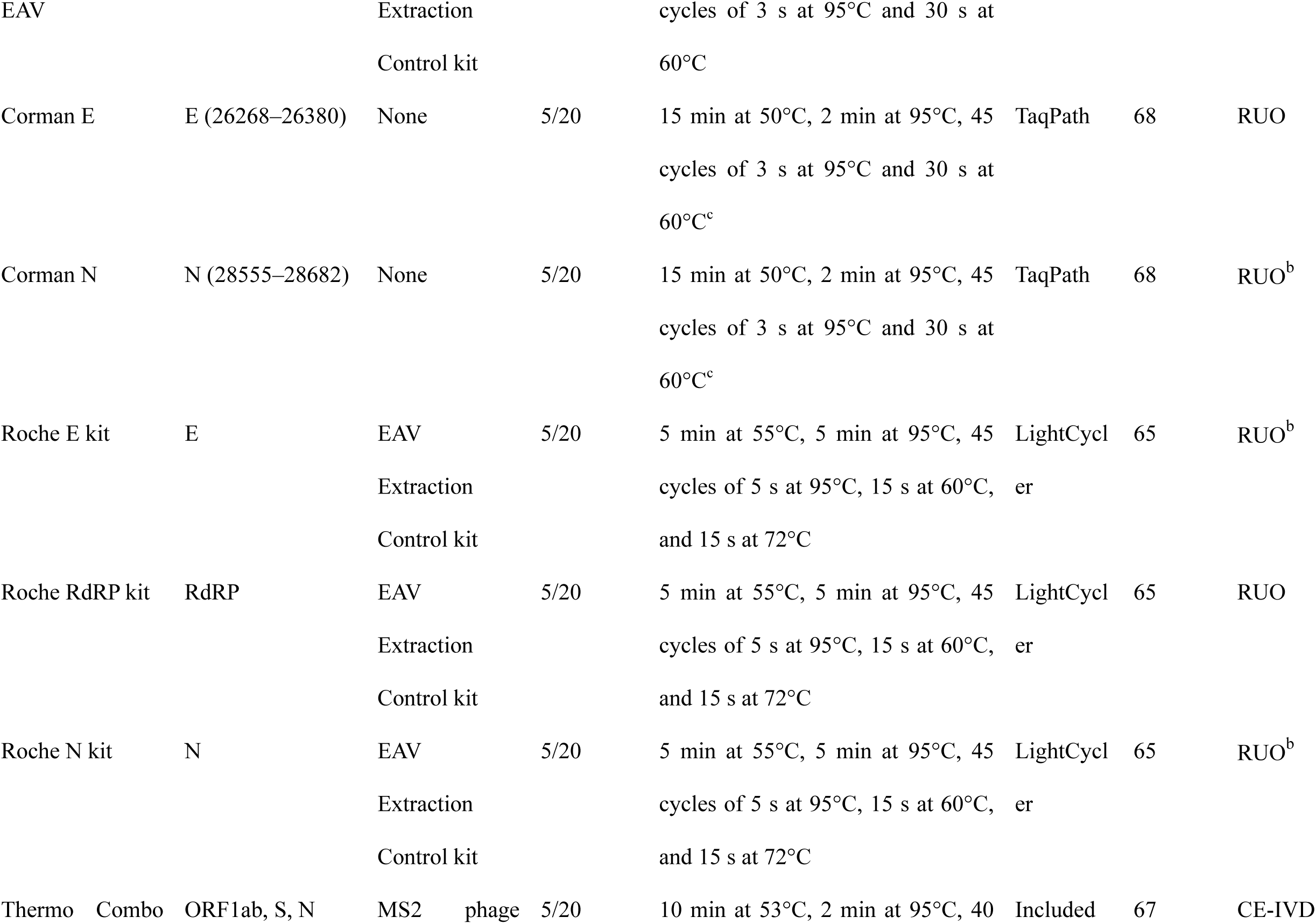
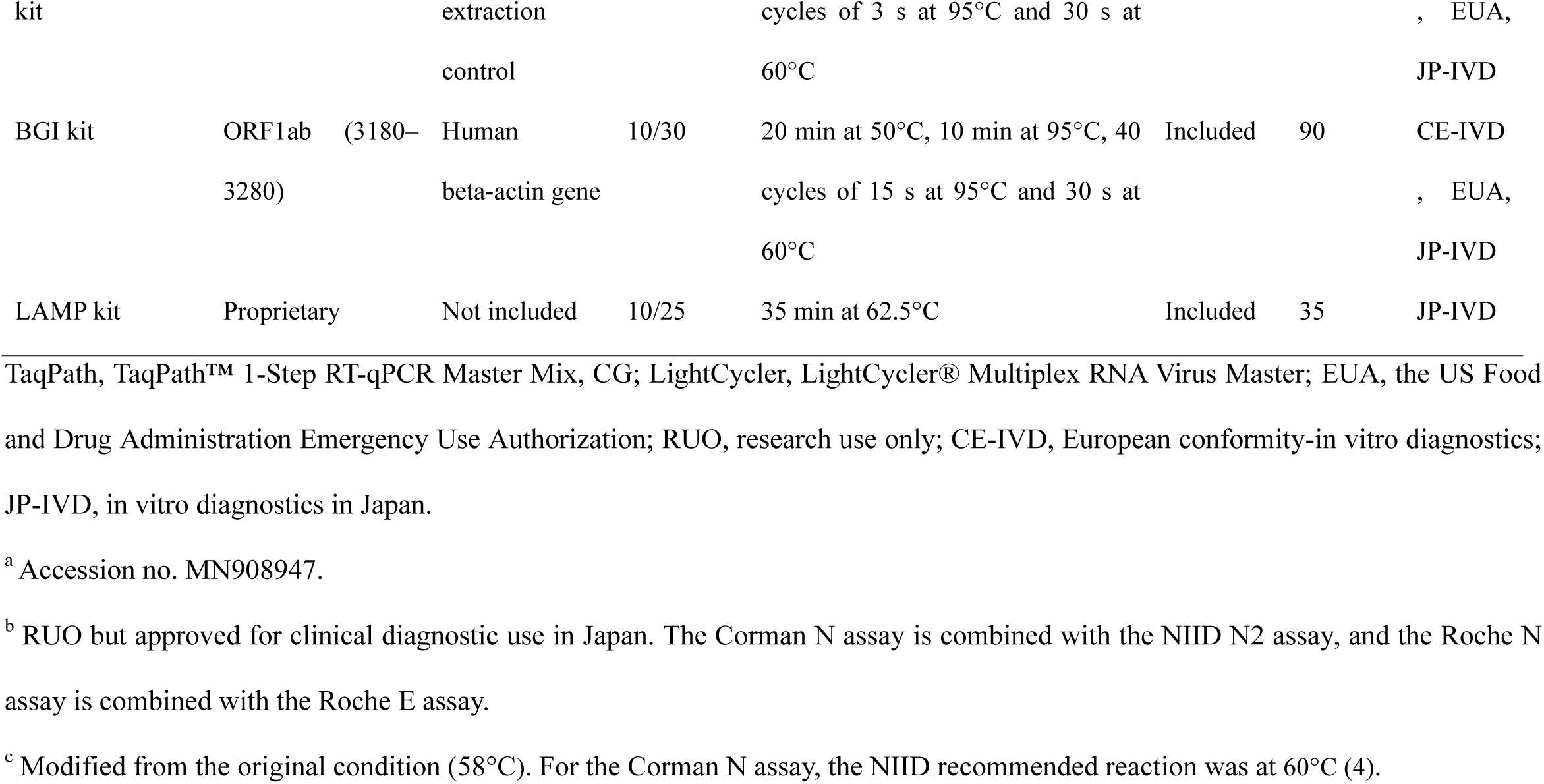
Summary of the molecular assays used in this study.

### Analytical sensitivity

We determined the limit of detection (LOD) of each assay using a minimum of four replicates of two-fold serial dilutions of recombinant Sindbis virus containing a partial SARS-CoV-2 genome (AccuPlex™ SARS-CoV-2 Reference Material Kit, 5,000 copies/mL; SeraCare, Milford, MA, USA). We calculated the 95% limit of detection (LOD) using probit analysis.

### Statistical analysis

At the time manuscript preparation, no gold standard exists. In this study, to ensure the presence of SARS-CoV-2 RNA and to avoid false-positives, a sample was defined as positive when positive test results were obtained for more than one genetic locus and assay and the others were defined as negative. The agreement of the assays was assessed by the Cohen’s kappa concordance coefficient. The sensitivity and specificity were compared using the McNemar test. The sensitivity of different specimen types was compared using the Fisher’s exact test. The Ct value were compared using the Kruskal– Wallis test or a Mann–Whitney U test. A *P*-value <0.05 was considered statistically significant. All statistical analyses were performed using SAS® Studio 3.8 (SAS Institute Inc., Cary, NC).

### Ethical statement

The Ethics Committee of Kyoto University Graduate School and the Faculty of Medicine approved this study (R2379).

## Results

A total of 155 study samples (65 nasopharyngeal swabs, 45 oropharyngeal swabs, and 45 sputum samples) were tested using the 12 assays. Sixty-eight samples (35 nasopharyngeal swabs, 15 oropharyngeal swabs, and 18 sputum samples) were positive for more than one assay and one genetic locus and were defined as true positive samples; the other samples were considered true negative. A full list of the results with Ct values is available as Dataset S1.

All the assays exhibited a specificity of 100%, while sensitivity varied (Table 2). The CDC N1, CDC N2, NIID N2 (with/without EAV), and Corman E assays were the most sensitive assays with ≥95.6% sensitivity. These 5 assays displayed high overall agreement compared with the reference standard (kappa values of ≥0.96) and between any two of them (kappa values of ≥0.95). The CDC N2 and NIID N2 assays exhibited 100% sensitivity; thus, their results were equal to the defined reference standard. The sensitivities of the remaining 7 assays (Corman N, Roche E, Roche RdRP, Roche N, Thermo Combo, BGI and LAMP assays; ≤88.2%) were significantly lower than those of the most sensitive assays.

**Table 2.**
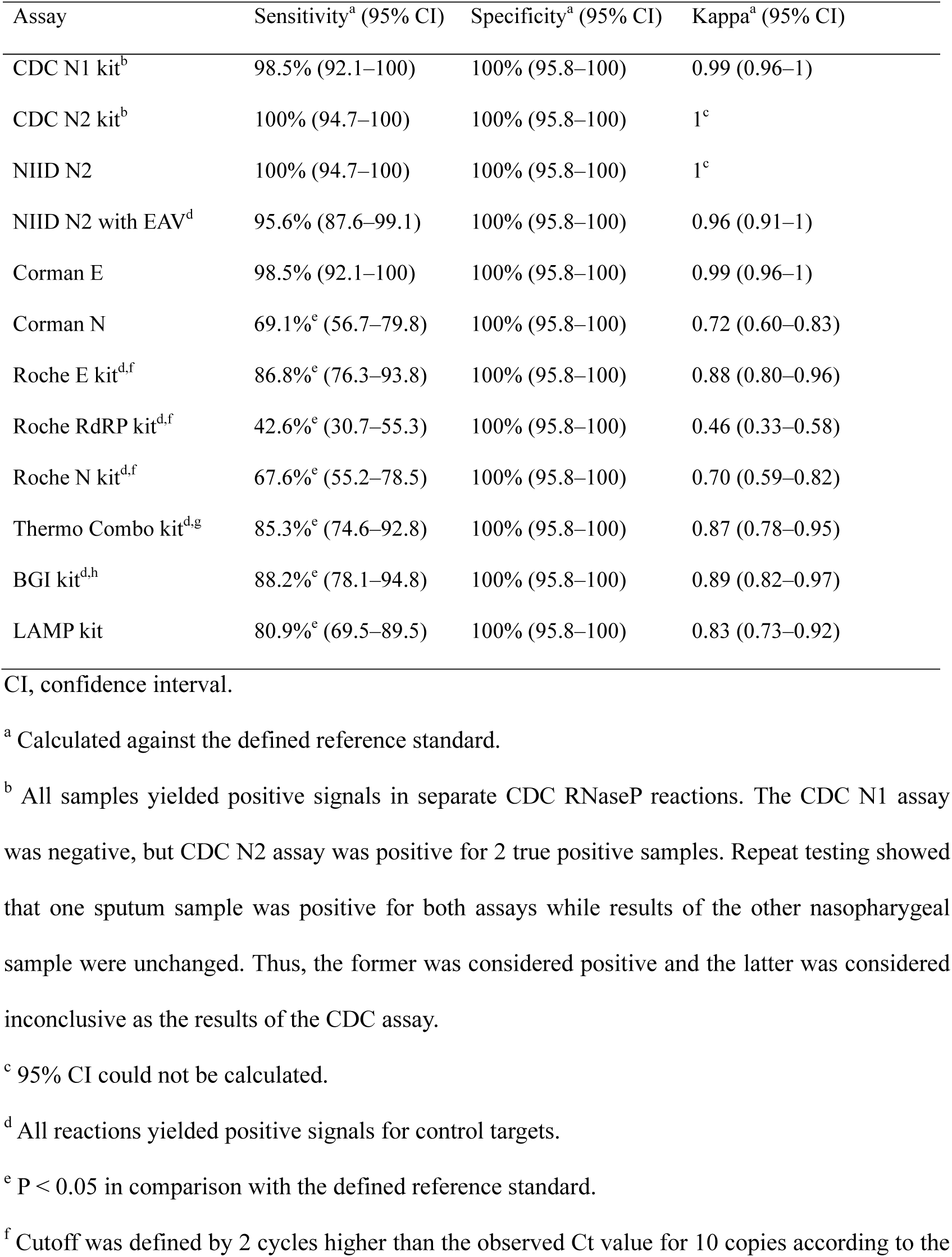
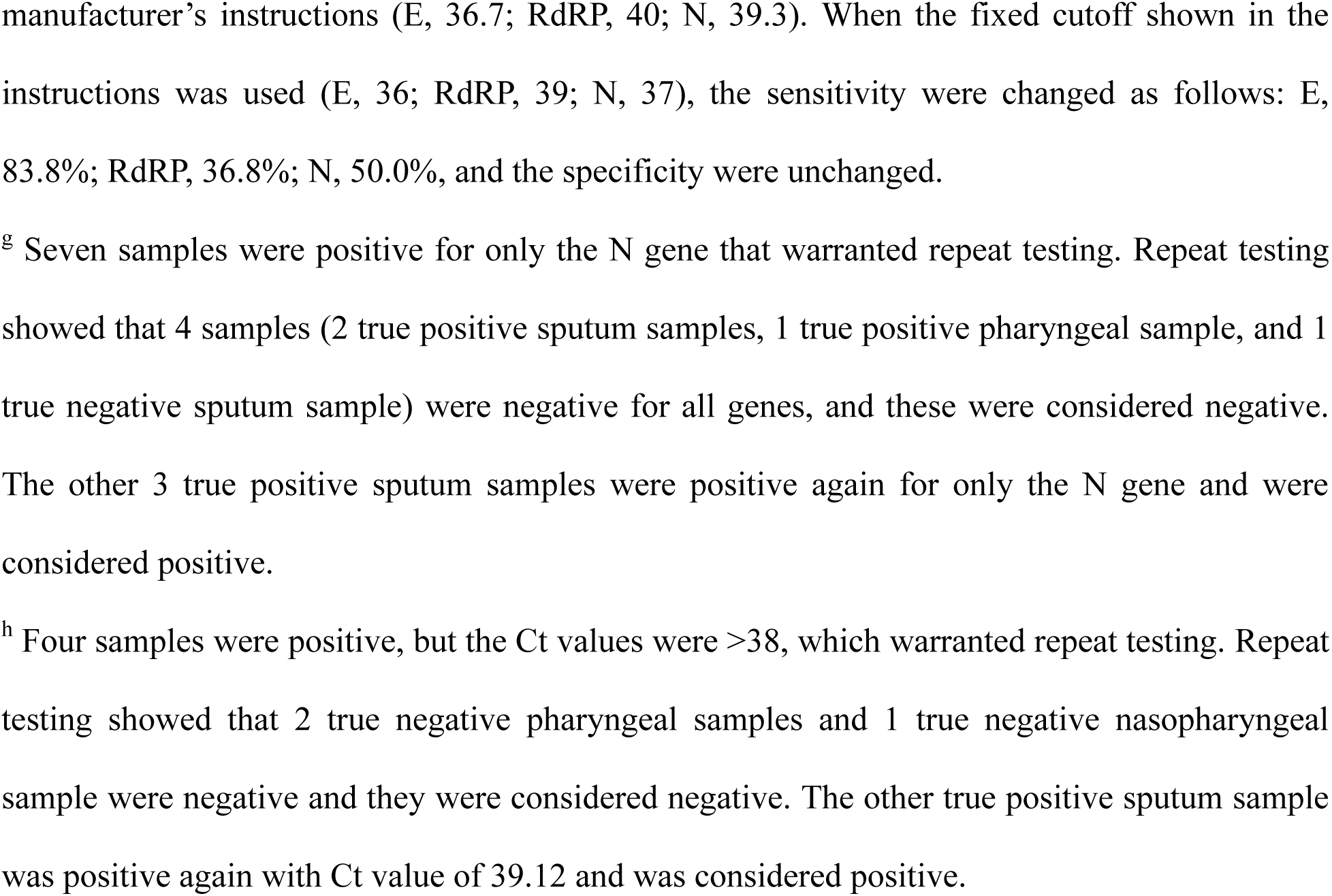
Overall diagnostic performance of 12 molecular assays.

| Assay | Sensitivity<sup>a</sup> (95% CI) | Specificity<sup>a</sup> (95% CI) | Kappa<sup>a</sup> (95% CI) |
| --- | --- | --- | --- |
| CDC N1 kit<sup>b</sup> | 98.5% (92.1–100) | 100% (95.8–100) | 0.99 (0.96–1) |
| CDC N2 kit<sup>b</sup> | 100% (94.7–100) | 100% (95.8–100) | 1<sup>c</sup> |
| NIID N2 | 100% (94.7–100) | 100% (95.8–100) | 1<sup>c</sup> |
| NIID N2 with EAV<sup>d</sup> | 95.6% (87.6–99.1) | 100% (95.8–100) | 0.96 (0.91–1) |
| Corman E | 98.5% (92.1–100) | 100% (95.8–100) | 0.99 (0.96–1) |
| Corman N | 69.1%<sup>e</sup> (56.7–79.8) | 100% (95.8–100) | 0.72 (0.60–0.83) |
| Roche E kit<sup>d,f</sup> | 86.8%<sup>e</sup> (76.3–93.8) | 100% (95.8–100) | 0.88 (0.80–0.96) |
| Roche RdRP kit<sup>d,f</sup> | 42.6%<sup>e</sup> (30.7–55.3) | 100% (95.8–100) | 0.46 (0.33–0.58) |
| Roche N kit<sup>d,f</sup> | 67.6%<sup>e</sup> (55.2–78.5) | 100% (95.8–100) | 0.70 (0.59–0.82) |
| Thermo Combo kit<sup>d,g</sup> | 85.3%<sup>e</sup> (74.6–92.8) | 100% (95.8–100) | 0.87 (0.78–0.95) |
| BGI kit<sup>d,h</sup> | 88.2%<sup>e</sup> (78.1–94.8) | 100% (95.8–100) | 0.89 (0.82–0.97) |
| LAMP kit | 80.9%<sup>e</sup> (69.5–89.5) | 100% (95.8–100) | 0.83 (0.73–0.92) |

CI, confidence interval.
<sup>a</sup> Calculated against the defined reference standard.
<sup>b</sup> All samples yielded positive signals in separate CDC RNaseP reactions. The CDC N1 assay was negative, but CDC N2 assay was positive for 2 true positive samples. Repeat testing showed that one sputum sample was positive for both assays while results of the other nasopharygeal sample were unchanged. Thus, the former was considered positive and the latter was considered inconclusive as the results of the CDC assay.
<sup>c</sup> 95% CI could not be calculated.
<sup>d</sup> All reactions yielded positive signals for control targets.
<sup>e</sup> $P < 0.05$ in comparison with the defined reference standard.
<sup>f</sup> Cutoff was defined by 2 cycles higher than the observed Ct value for 10 copies according to the
manufacturer's instructions (E, 36.7; RdRP, 40; N, 39.3). When the fixed cutoff shown in the instructions was used (E, 36; RdRP, 39; N, 37), the sensitivity were changed as follows: E, 83.8%; RdRP, 36.8%; N, 50.0%, and the specificity were unchanged.
<sup>g</sup> Seven samples were positive for only the N gene that warranted repeat testing. Repeat testing showed that 4 samples (2 true positive sputum samples, 1 true positive pharyngeal sample, and 1 true negative sputum sample) were negative for all genes, and these were considered negative. The other 3 true positive sputum samples were positive again for only the N gene and were considered positive.
<sup>h</sup> Four samples were positive, but the Ct values were >38, which warranted repeat testing. Repeat testing showed that 2 true negative pharyngeal samples and 1 true negative nasopharyngeal sample were negative and they were considered negative. The other true positive sputum sample was positive again with Ct value of 39.12 and was considered positive.

The CDC protocol requires both N1 and N2 assays, and a sample will be considered positive if both produced positive results. In this study, one true positive nasopharygeal sample was positive only for the N2 assay even after retesting. The sample was considered inconclusive and the performance of the CDC protocol was considered the same as the CDC N1 assay. The NIID protocol includes both NIID N2 and Corman N assays, and a sample will be considered positive if either assay produces a positive result. In this study, 69.1% of samples were positive for both assays, and 30.9% were positive for only the N2 assay. The protocol by Corman recommended an E assay that detects SARS-related viruses (*Sarbecoviru*s) as a first-line screening assay and then SARS-CoV-2 specific RdRP assay for confirmatory testing (1). This approach defined only 49.2% of the Roche E assay-positive samples as SARS-CoV-2, although a single positive result of the Corman E or Roche E assay can be interpreted as SARS-CoV-2 positive in the absence of other *Sarbecovirus*. Assays with multiplexed internal control reactions and the CDC RNaseP assay yielded positive signals for all samples.

Table 3 shows diagnostic performances for each specimen type. Nasopharyngeal swabs tended to have a higher sensitivity than the other samples. The sensitivity of Corman N assay for sputum samples and that of Roche N assay for oropharyngeal swabs and sputum samples were significantly lower than those for nasopharyngeal swabs. The Ct values of CDC N2 and NIID N2 assays for nasopharyngeal swabs (median [interquartile range], 27.1 [23.6–31.1] and 29.7 [26.3–33.3], respectively) were lower than those for oropharyngeal swabs (31.5 [29.9–35.0] and 33.0 [32.0–34.6]) or sputum samples (30.0 [25.6–33.5] and 30.9 [28.3–34.0]) but the differences did not reach statistical significance (P=0.11 and 0.16 by comparison among 3 specimen types, respectively). The sputum samples had the lower Ct values of the CDC RNaseP assay (25.6 [23.6–27.8]) than nasopharyngeal or oropharyngeal swabs (28.2 [26.9 –29.6], P<0.001 and 28.8 [26.9–31.0], P<0.001, respectively).

**Table 3.**
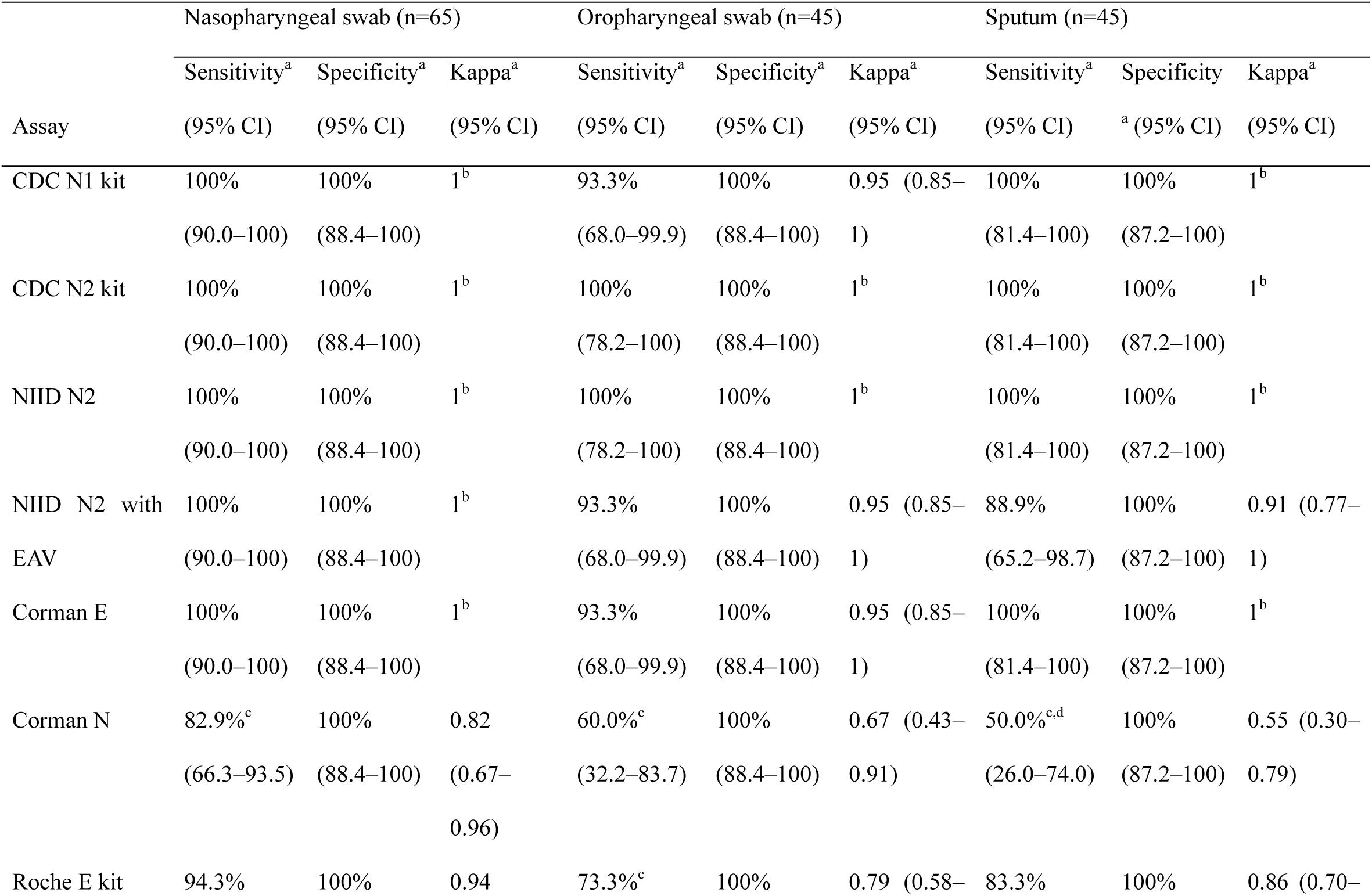
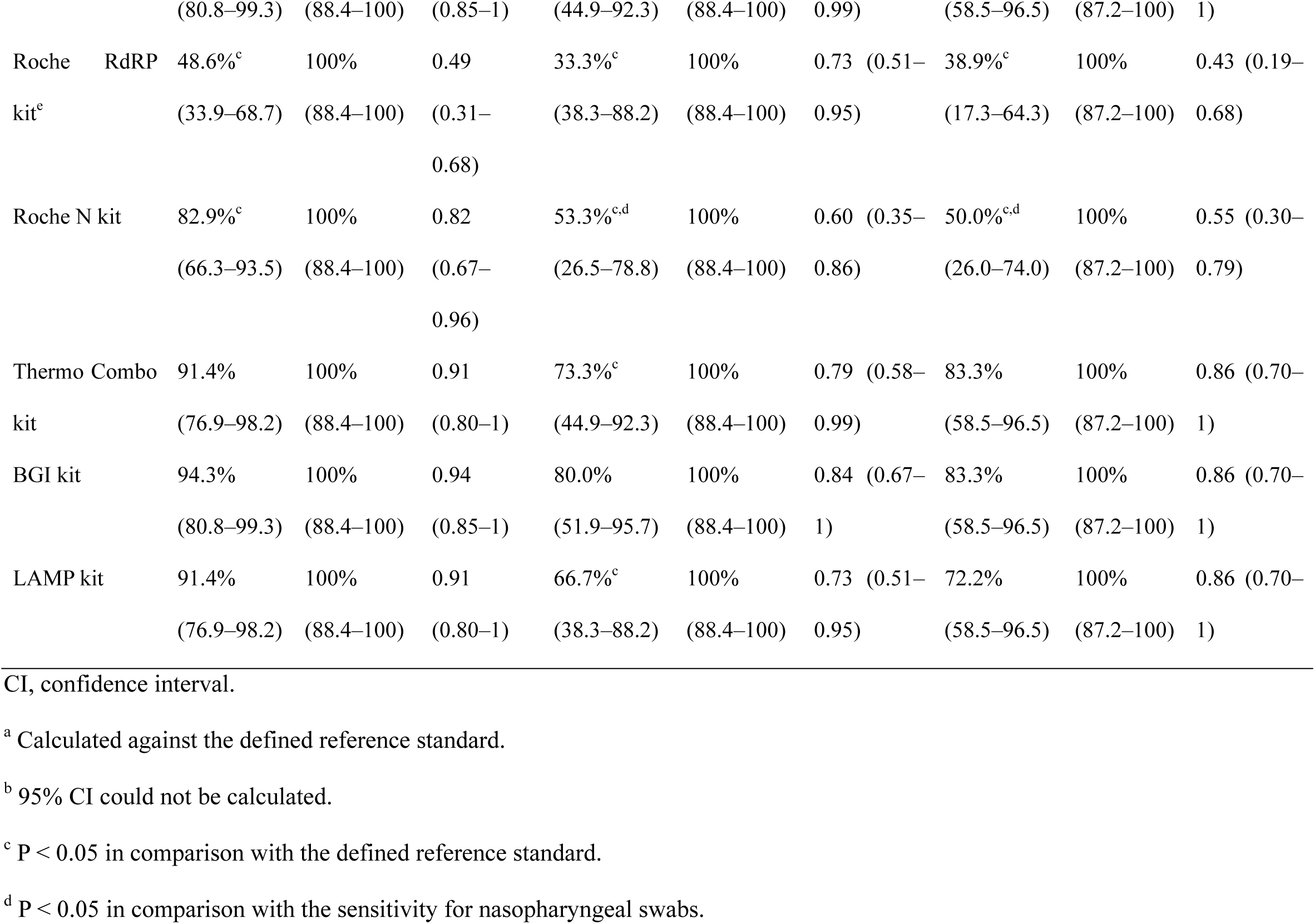
Diagnostic performance of 12 molecular assays according to specimen types.

The lowest LOD was observed for the CDC N2 assay (136 copies/mL or 1.6 copies/reaction). The other assays which showed the high sensitivities in clinical samples (CDC N1, NIID N2, and Corman E assays) displayed LODs of 191–271 copies/mL. The LODs of the Roche RdRP and N assays were high (>5,000 copies/mL).

## Discussion

The current diagnosis of COVID-19 mainly relies on RT-PCR tests (6). We performed manufacturer-independent evaluation of the molecular assays, including commercial kits that utilize otherwise-extracted RNA templates. We found that the specificity was perfect for all the assays and that the CDC N1, CDC N2, NIID N2, and Corman E assays were the most sensitive and highly concordant (7). Genetic variations that may compromise sensitivity of the CDC N1, N2, and Corman E assays have been rarely observed as of week 21 of 2020 (8). False negatives by the other assays occurred among low-copy number samples (presenting high Ct values by the CDC N2 or NIID N2 assay; Dataset S1), suggesting a lack of sensitivity of these assays.

The Roche assays were based on Corman’s assays (1) but had lower sensitivity for their E and N assays. This is likely due to lower Ct cutoffs for the Roche assays, rather than differences in reagents and reaction conditions (Table 1 and Dataset S1). Previous studies reported that N assay was less sensitive than the E and RdRP assays (1) and the RdRP assay was less sensitive than the Roche E assay (3). The low sensitivity of the Roche RdRP and N assays were concordant with their high LODs (Table 4). The low sensitivity of the BGI assay may be due to the inclusion of human gene internal controls in the same reaction, which could prevent amplification of viral genes, especially in human genome-enriched samples. The LAMP assay can be used in a resource-poor setting and has the fastest assay time due to its isothermal reaction. However, it had a low sensitivity and no control reaction. The lower sensitivities observed for oropharyngeal and/or sputum samples might be related to higher viral copies (lower Ct values) in nasopharyngeal swabs and/or higher copies of human genes (lower Ct values of the CDC RNaseP Assay) in sputum samples.

**Table 4.**
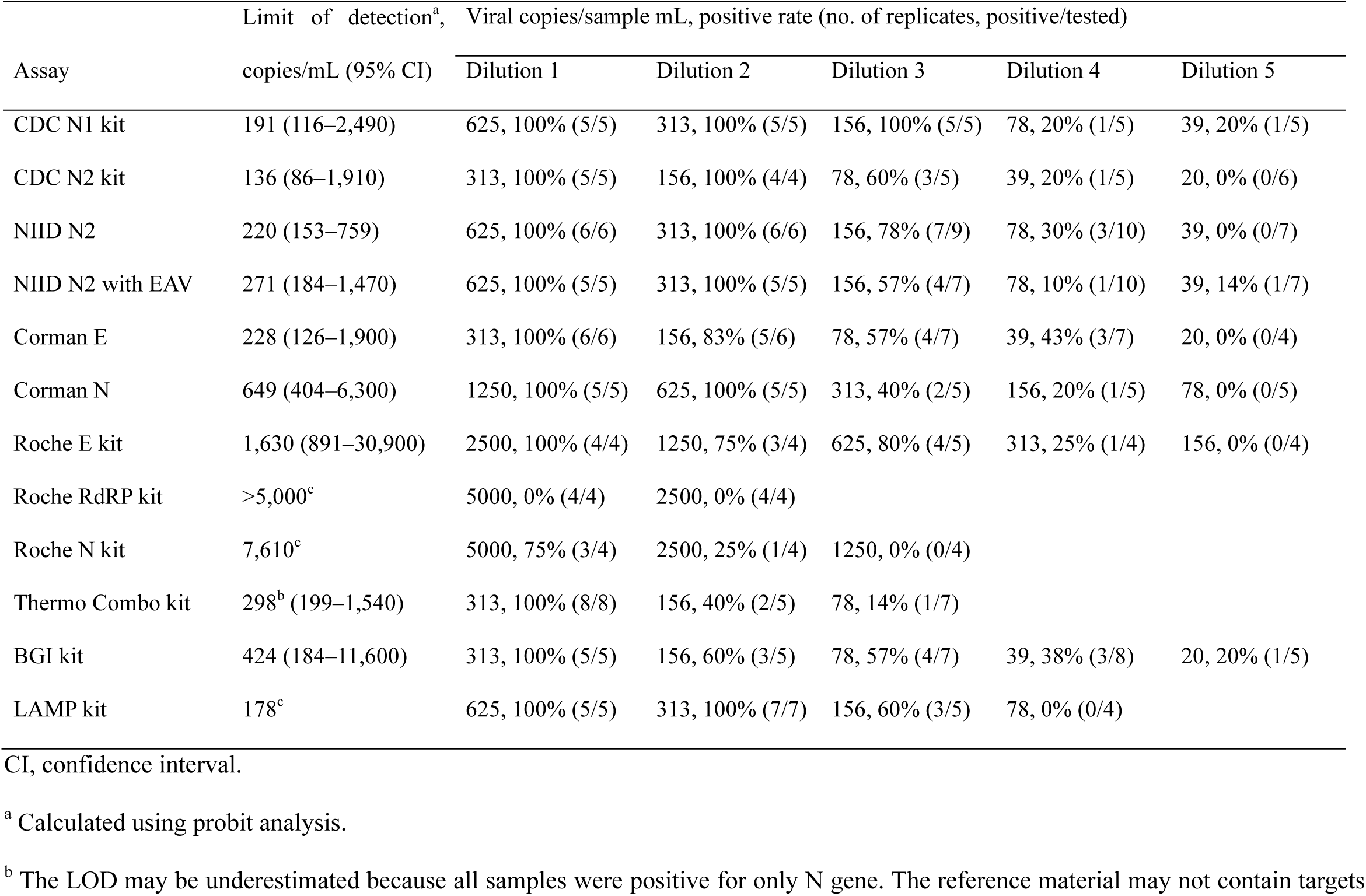
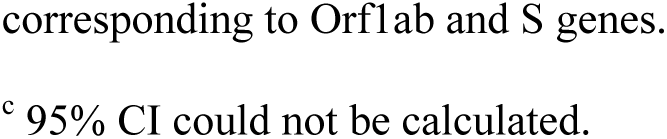
Analytical sensitivity of 12 molecular assays.

| Assay | Limit of detection<sup>a</sup>,<br>copies/mL (95% CI) | Viral copies/sample mL, positive rate (no. of replicates, positive/tested) |  |  |  |  |
| --- | --- | --- | --- | --- | --- | --- |
|  |  | Dilution 1 | Dilution 2 | Dilution 3 | Dilution 4 | Dilution 5 |
| CDC N1 kit | 191 (116–2,490) | 625, 100% (5/5) | 313, 100% (5/5) | 156, 100% (5/5) | 78, 20% (1/5) | 39, 20% (1/5) |
| CDC N2 kit | 136 (86–1,910) | 313, 100% (5/5) | 156, 100% (4/4) | 78, 60% (3/5) | 39, 20% (1/5) | 20, 0% (0/6) |
| NIID N2 | 220 (153–759) | 625, 100% (6/6) | 313, 100% (6/6) | 156, 78% (7/9) | 78, 30% (3/10) | 39, 0% (0/7) |
| NIID N2 with EAV | 271 (184–1,470) | 625, 100% (5/5) | 313, 100% (5/5) | 156, 57% (4/7) | 78, 10% (1/10) | 39, 14% (1/7) |
| Corman E | 228 (126–1,900) | 313, 100% (6/6) | 156, 83% (5/6) | 78, 57% (4/7) | 39, 43% (3/7) | 20, 0% (0/4) |
| Corman N | 649 (404–6,300) | 1250, 100% (5/5) | 625, 100% (5/5) | 313, 40% (2/5) | 156, 20% (1/5) | 78, 0% (0/5) |
| Roche E kit | 1,630 (891–30,900) | 2500, 100% (4/4) | 1250, 75% (3/4) | 625, 80% (4/5) | 313, 25% (1/4) | 156, 0% (0/4) |
| Roche RdRP kit | >5,000<sup>c</sup> | 5000, 0% (4/4) | 2500, 0% (4/4) |  |  |  |
| Roche N kit | 7,610<sup>c</sup> | 5000, 75% (3/4) | 2500, 25% (1/4) | 1250, 0% (0/4) |  |  |
| Thermo Combo kit | 298<sup>b</sup> (199–1,540) | 313, 100% (8/8) | 156, 40% (2/5) | 78, 14% (1/7) |  |  |
| BGI kit | 424 (184–11,600) | 313, 100% (5/5) | 156, 60% (3/5) | 78, 57% (4/7) | 39, 38% (3/8) | 20, 20% (1/5) |
| LAMP kit | 178<sup>c</sup> | 625, 100% (5/5) | 313, 100% (7/7) | 156, 60% (3/5) | 78, 0% (0/4) |  |

CI, confidence interval.
<sup>a</sup> Calculated using probit analysis.
<sup>b</sup> The LOD may be underestimated because all samples were positive for only N gene. The reference material may not contain targets
corresponding to Orf1ab and S genes.
<sup>c</sup> 95% CI could not be calculated.

To avoid false negatives due to technical errors such as extraction problems or PCR inhibition, it is recommended to include internal control reactions. The CDC assays were designed to be combined with a separate internal control reaction (Table 1). Different from multiplex assays that incorporate internal controls such as Roche, Thermo, or BGI kits, this approach needs extra reagents, time, and space in a reaction plate but can be combined with other in-house assays (NIID N2 or Corman E) without any modification. For the multiplex approach, we selected the NIID N2 assay to be multiplexed with the Roche EAV kit, resulting in the similar performance as the original NIID N2 assay.

To date, two published reports have compared performances of multiple RT-PCR assays using clinical samples. Nalla (9) et al. compared CDC N1/N2/N3 and Corman E/RdRP among 10 SARS-CoV-2-positive samples. They reported that the CDC N2 and Corman E assays were the most sensitive. van Kasteren et al. compared 7 commercial kits, including 13 positive and 6 negative samples (10). When compared with the Corman E assay, the R-Biopharm AG performed the best, followed by BGI, KH Medical, and Seegene. These reports are in agreement with our findings.

The study limitations included a relatively small sample size of each specimen type and lack of clinical information, measurements by multiple investigators, and genomic variation analysis.

In conclusion, we validated the NIID N2 assay with EAV control reaction and found that the CDC EUA kit (N1/N2/RNaseP), NIID N2 with/without EAV, and Corman E assays were the most sensitive assays. They are feasible as references and clinical diagnostic tests until commercial kits with internal control reactions or fully automated systems that have high diagnostic performances are available in clinical laboratories. Continuous efforts to improve COVID-19 diagnostics are important to control this pandemic.

## Acknowledgments

This research received no specific grant from any funding agency in the public, commercial, or not-for-profit sectors. We have no conflict of interest to declare. We thank Akihiko Matsuo (Department of Clinical Laboratory, Kyoto University Hospital, Kyoto, Japan) and Akihiko Hayashi (Department of Clinical Laboratory, Kyoto City Hospital, Kyoto, Japan) for their technical assistance.

